# Comparing Livestock Mobility-Informed Strategies for Peste des Petits Ruminants Control in Nigeria: The Central Role of the Network Backbone

**DOI:** 10.64898/2025.12.16.693877

**Authors:** Asma Mesdour, Sandra Ijoma, Muhammad-Bashir Bolajoko, Mamadou Ciss, Stephen Eubank, Eric Cardinale, Mathieu Andraud, Andrea Apolloni

## Abstract

Animal mobility is central to pastoral livelihoods and regional trade in West Africa, but it also facilitates the spread of transboundary animal diseases such as Peste des petits ruminants (PPR). In Nigeria, PPR outbreaks recur regularly, yet surveillance and control remain limited in the absence of routine animal-movement tracking. Here, we assess and compare movement-informed control options for PPR using a reconstructed livestock mobility network from a one-time market survey conducted in three northern Nigerian states. We simulate transmission on this network and evaluate three intervention strategies: (i) targeting vulnerable villages, (ii) targeting the links that connect movement communities, and (iii) targeting villages belonging to the network backbone. Across scenarios, backbone-based targeting consistently produced the largest reductions in network connectivity and epidemic outcomes, outperforming strategies focused on vulnerable nodes or inter-community links.

These results suggest that backbone-informed control could provide a practical, resource-efficient pathway to strengthen PPR control in settings where routine movement data are scarce.

## Introduction

Transboundary animal diseases (TADs) are highly contagious infections that can spread rapidly across borders, generating major socioeconomic risks. Among these, peste des petits ruminants (PPR) is a severe morbilliviral disease of domestic small ruminants (goats and sheep), often associated with high morbidity and mortality (Njeumi et al. 2020). First reported in 1942 (initially described in Côte d’Ivoire), PPR expanded across much of Africa, the Middle East, and Asia (Banyard et al. 2010). After decades of absence from the European Union, new incursions were detected in 2024–2025, with outbreaks reported in Greece and Romania (July 2024), followed by Bulgaria (late 2024) and Hungary (early 2025) (Parida et al. 2024; Imanbayeva et al. 2025).

Globally, small ruminants number ∼2.1 billion head, and ∼80% are concentrated in Africa and Asia, regions where PPR remains a major constraint on food security and poverty reduction. Recognizing this burden, the Food and Agriculture Organization (FAO) and the World Organisation for Animal Health (WOAH) set the goal of global PPR eradication by 2030, using a stepwise strategy in which strengthened surveillance and optimized vaccination are foundational (Njeumi et al. 2020; Imanbayeva et al. 2025). However, translating this roadmap into impact requires country-specific operationalization. In many endemic settings, weak infrastructure, limited cold-chain capacity, and constrained vaccination logistics continue to impede consistent implementation (Wane et al. 2020; Chukwudi et al. 2020).

Because livestock mobility connects otherwise separated populations, animal movements can accelerate PPR dissemination and complicate eradication efforts. Consequently, network analysis of movement systems has become a key approach for quantifying potential transmission pathways and identifying leverage points for targeted surveillance and control (Nicolas et al. 2018; Apolloni et al. 2019). In West Africa, market- and transhumance-linked movement networks are typically highly heterogeneous, with a small number of highly connected “hub” locations and a subset of “bridging” connections that link communities (clusters). In principle, interventions such as risk-based vaccination focused on hubs, time-bounded measures around high-connectivity markets, and selective management of bridging connections could reduce spread disproportionately, aligning well with the FAO/WOAH emphasis on smarter control (e.g., vaccination under resource constraints) (FAO, 2015; Gates and Woolhouse 2015; Dione et al., 2025). Because of factors related to the commercial chain, social interactions, and resource availability, some locations form tightly connected communities of animal movements, which are linked to others mainly through a limited number of bridging connections (Jahel et al. 2020).

Yet key gaps remain: many movement-network studies in the region rely on short observation windows and partial spatial coverage, using ad hoc market surveys, interviews, or compiled survey datasets rather than continuous animal identification and movement traceability systems (Baazizi et al. 2017; Apolloni et al. 2019).

Nigeria is a regional demographic and economic giant, with a 2025 population estimate of ∼237.5 million. Agriculture remains an important contributor to the economy (e.g., ∼22.7% of GDP in 2023), and the livestock sector is frequently cited as contributing ∼5% of national GDP and ∼17% of agricultural GDP. Recent national reporting also underscores the very large scale of small-ruminant holdings. PPR has been reported in Nigeria for decades, and the absence of routine movement traceability hinders timely identification of risk corridors and prioritization of surveillance and vaccination (Ijoma et al. 2025). Recent work has begun to address this by reconstructing small-ruminant mobility networks in three northern states using mixed survey and participatory methods and by using simulation/network metrics to identify vulnerable or sentinel areas (Mesdour et al. 2024; Ijoma et al. 2025). However, the extent to which these “mobility-defined hotspots” are also the most efficient operational targets for control (e.g., targeted vaccination vs. disruption of bridging links) remains uncertain.

Here, we test whether mobility-defined surveillance hotspots, high-centrality nodes, bridging links, and network backbones are also the most efficient targets for control. Focusing on three northern Nigerian states, we investigate PPR transmission dynamics and compare alternative strategies, including targeted removal or protection of vulnerable nodes, inter-cluster links, and the backbone. By leveraging network analysis under realistic operational constraints, our goal is to provide actionable evidence to enhance PPR control in regions at highest risk.

## Material & Methods

### Data and Small Ruminant Movement Network Reconstruction

Based on a preliminary analysis conducted by partners of the Livestock Disease Surveillance Knowledge Integration (LIDISKI) project ^1^, ten markets in three Nigerian states were selected for sampling. Six markets were sampled in Plateau, and for security, logistical, and accessibility reasons, only two markets were sampled in both Bauchi and Kano States (Ijoma et al. 2025). Data on origin and destination, and the number of small ruminants moved, were collected. From these data, we reconstructed the small ruminant’s mobility network whose nodes are villages, and directed links represent movements from origin to destination, weighted by the number of animals moved.

### Identification of Contagion Clusters Based on Disease Dynamics Within the Small Ruminant Movement Network

The identification of contagion clusters or communities, areas with strong interconnectivity and shared vulnerability, can further refine control efforts. These clusters, characterized by their potential for rapid disease spread, underscore the need for targeted interventions (Salathé and Jones 2010; Nath et al. 2019; Mishra et al. 2023).

This step follows the COCLEA algorithm (COntagion CLusters Extraction Algorithm) (Nath et al. 2019) to identify contagion clusters (CC), i.e., sets of nodes (villages) that share a common vulnerability to contagion. An infection occurring at a node *i* ∈ CC1 substantially increases the probability that other nodes *j* ∈ CC1 will also be infected. While this concept aligns with the idea of communities in a network, it places a stronger focus on the transmission dynamics of the infection across the network structure (Nath et al. 2019). This algorithm identifies such groups through two main steps:

- Link Ranking: Each link is assigned a score reflecting its importance in the spread of infection. This is estimated by analyzing how often the link appears in critical transmission paths across random subgraphs of the network.
- Link Removal: Links are then removed in order of decreasing importance until the largest strongly connected component (SCC) is reduced below a specified size *n*.

Following the approach of Nath (2019), we varied the *n* size, which corresponds to the desired number of nodes in the SCC, using values *n*=2, 5, 10, 20, 30, and |V|.V corresponds to the total number of nodes at the village level. We then compared the number of detected contagion clusters and their distribution areas. Based on this comparison, the optimal size of *n* was selected by choosing the best balance, determined as the most appropriate compromise between these *n*. In particular, the *n* encompasses a broad range of areas.

### Backbone Extraction Using a Weighted Small Ruminant Movement Network

Enhancing control hinges on identifying critical areas and dominant transmission pathways. One promising approach involves isolating a network subset that represents regions with the highest density of animal movement, commonly referred to as the “backbone” (Neal 2022; Correia et al. 2023). This backbone is vital for simplifying complex networks, enabling researchers to uncover underlying dynamics while filtering out sporadic or inconsequential interactions that contribute minimally to the phenomenon under scrutiny (Ferreira et al. 2022). By focusing on the network backbone, it is possible to highlight key movement patterns, thereby increasing the potential of a strategic area to monitor. This backbone not only highlights critical mobility structures but also provides a valuable framework for designing effective surveillance approaches by targeting the areas forming the backbone (Mesdour et al. 2024). However, its influence on epidemic control outcomes remains largely unexplored. In this step, this backbone was isolated; The small ruminants mobility network in this study, as mentioned previously, is weighted by the number of animals moving between nodes, where the edges are characterized by weights representing the strength of relationships between nodes. We used the disparity filter method (Serrano et al. 2009) to filter links. This algorithm is based on the *p-value* statistical significance of the null model, i.e., where for each node, the weights are distributed randomly among the links. Using the disparity filter at significance level *α* = 0.07, we computed for each edge *(i, j)* the *p-value α*_*ij*_, the probability, under the null, of observing a normalized weight ≥ *w*_*ij*_. Edges with *α*_*ij*_ *≤ α* were retained; all others were pruned. We implemented this with the backbone package (Neal 2022).

### Impact of the Target Control on the Epidemic Simulation Outcomes

In this section, we explored various scenarios that could benefit from improving the control of PPR (Table 1). Using an SIR (Susceptible, Infectious, Recovered) model, we simulated control scenarios by removing nodes and/or links. The underlying hypothesis is that if controls are focused on these nodes, disease spread will be limited from these critical points.

**Table 1.** Six simulated intervention scenarios (S1–S6): vulnerable-node removal by severity (Q1–Q4), cluster-bridge closures, and backbone targeting.

| Scenario | Intervention Approach | Description |
| --- | --- | --- |
| 1 | Remove Q1 vulnerable-node set | Upon first case detection, villages in the Q1 set are closed to movement (entry/exit bans) to contain the spread. |
| 2 | Remove Q2 vulnerable-node set | Upon first case detection, villages in the Q2 set are closed to movement (entry/exit bans). |
| 3 | Remove Q3 vulnerable-node set | Upon first case detection, villages in the Q3 set are closed to movement (entry/exit bans). |
| 4 | Remove Q4 vulnerable-node set | Upon first case detection, villages in the Q4 set are closed to movement (entry/exit bans). |
| 5 | Cut interlinks between contagion clusters | Upon first case detection, roads/edges that bridge identified contagion clusters are immediately closed. |
| 6 | Target backbone villages | Upon first case detection, villages on the network backbone receive movement bans (entry/exit) and vaccination. |

Previously, using simulations on the small-ruminant movement network, we identified as sentinel candidates the nodes most often infected before the epidemic peak (Mesdour et al. 2024). Different sets of vulnerable nodes were identified as a function of epidemic severity. In the underlying simulation framework, epidemic outcomes were generated across combinations of the per-movement transmission probability (*Pinf*) and the probability of recovery (*Prec*). Simulated epidemics were then classified according to their resulting epidemic size, yielding four severity groups (Q1–Q4) ranging from low- to high-severity outbreaks. Thus, Q1–Q4 do not represent clinical severity categories but rather classes of simulated epidemic severity derived from the distribution of epidemic outcomes across parameter combinations. These groups were used here to evaluate whether intervention performance remained consistent across a gradient of epidemic conditions.

For each severity group (Scenarios 1–4), control measures were applied to the corresponding set of vulnerable (sentinel) nodes identified for that epidemic severity level. As the number and identity of sentinel nodes differ among Q1–Q4 groups, these scenarios enabled us to assess whether intervention effectiveness varied across a gradient of epidemic severity (Table 1).

Furthermore, we identified the ‘bridges’ connecting contagion clusters, which are considered Scenario 5 (Table 1). These bridges represent critical links between contagion clusters within the network. We hypothesized that targeting control efforts on these bridges could be effective and therefore tested the impact of removing these connections; If a disease is declared in one of the contagion clusters, we can forbid or restrict the movement within these links that connect the contagion clusters, and this could stop the spread of the disease to a group of villages.

Another scenario tested involves the removal of backbone nodes in the network, referred to as Scenario 6. Since this structure is designed to retain only the most important information, targeting it through vaccination and/or movement restrictions could be an effective control strategy (Table 1).

To compare scenarios, we evaluate three network metrics, the Largest Connected Component (LCC), Efficiency (Eff), and Total Flow (TF), on the networks obtained after each intervention. Together, these indicators summarize changes in structure, performance, and resilience. The LCC (the “giant component”) is the size of the largest set of mutually reachable nodes in the network (Bellingeri et al. 2020). In weighted networks, efficiency is computed from shortest-path distances derived from edge weights (Latora and Marchiori 2007). Higher (global) efficiency means nodes can reach each other in fewer steps on average; a drop in efficiency indicates longer routes and more unreachable pairs (counted as zero), reflecting a less well-connected network (Bellingeri et al. 2020). Total Flow (TF) is the overall quantity of animals moving through the connections or pathways within the network.

Finally, we simulated transmission to quantify each scenario’s effect on the final epidemic size, comparing results to a no-control reference network.

## Results

### Small Ruminants Mobility Network Description

Details concerning the network of small ruminant mobility in the northern state of Nigeria are detailed in Ijoma et al. (2025) (Ijoma et al. 2025). After aggregating movements at the village level, the resulting network comprised 235 nodes (villages) and 355 weighted links, where link weights represent the number of small ruminants moved between villages. Of the 355 recorded movements, 171 (48.17%) occurred entirely within the Lidiski study area (Plateau, Bauchi, and Kano). A further 136 movements (38.31%) originated within Lidiski but were destined for locations outside the study area, whereas only 9 movements (2.54%) entered Lidiski from outside (i.e., had destinations within Lidiski but origins elsewhere). The remaining movements occurred entirely outside the Lidiski area but were retained because they connect sampled origins/destinations in the reconstructed network.

Geographically, inter-village distances spanned from approximately 6 km to 922 km (mean: 361 km).

Contagion clusters were geographically dispersed across Nigeria. Using COCLEA, we detected similar patterns of dispersed clusters across all tested values of *n*. We retained *n* = 20 as a compromise between smaller settings (*n* = 10 and *n* = 15); importantly, some southern nodes that drop out at *n* = 15 remain included at *n* = 20. With this choice, the network partitions into seven communities spanning 17 states across both northern and southern Nigeria (Figure 1).

**Figure 1.**
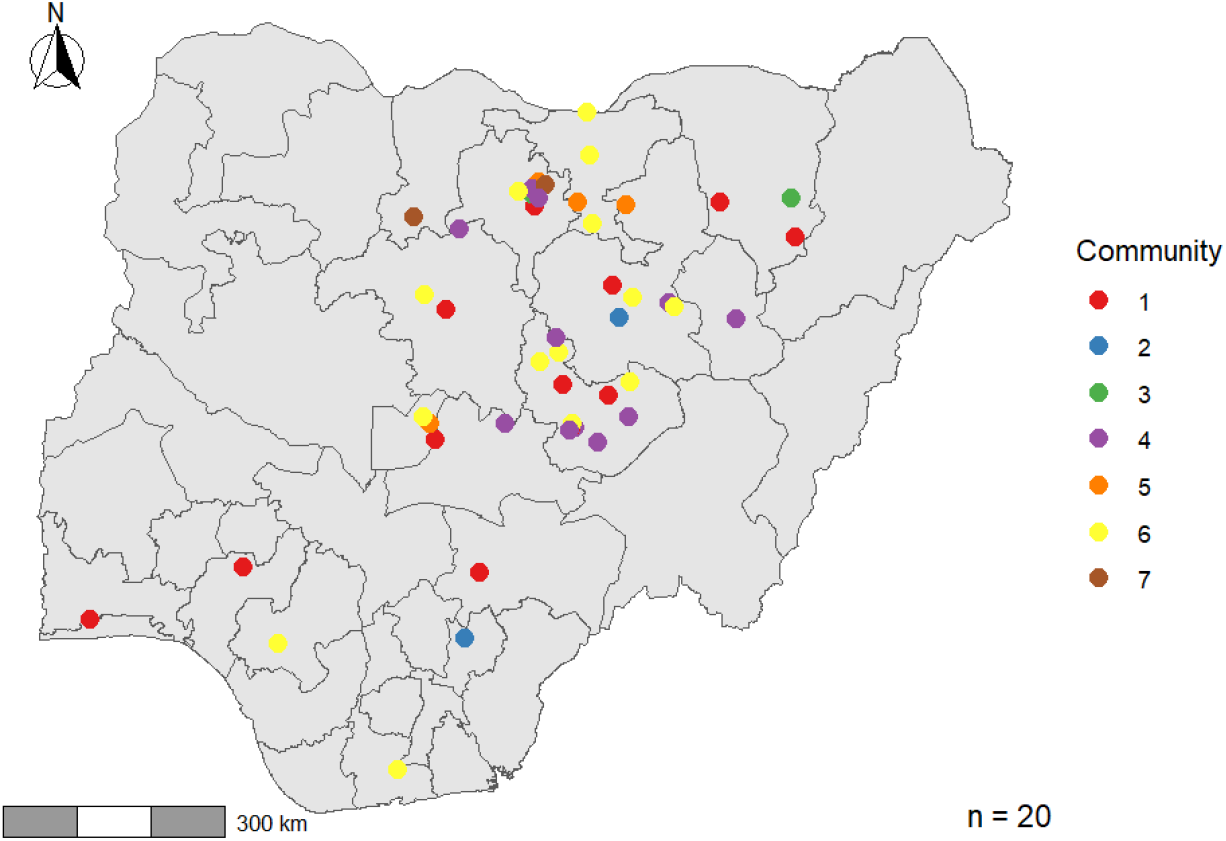
Contagion Clusters in the Small Ruminants Mobility Network. The map illustrates the geographic distribution of contagion clusters within the network (identified using the COCLEA algorithm (COntagion CLusters Extraction Algorithm), based on a target strongly connected component (SCC) size of n = 20. Colors indicate the clusters, revealing seven distinct contagion clusters. These clusters are geographically dispersed. Nodes not belonging to any of the identified clusters are not displayed.

The largest community contains 16 nodes distributed across eight states (Bauchi, Edo, the Federal Capital Territory, Jigawa, Kaduna, Kano, Plateau, and Rivers). It is followed by communities of 15 nodes across seven states (Bauchi, Gombe, Kaduna, Kano, Nasarawa, Plateau, and Yobe), 13 nodes across nine states (Bauchi, Benue, the Federal Capital Territory, Kaduna, Kano, Lagos, Ondo, Plateau, and Yobe), 6 nodes across four states (Bauchi, the Federal Capital Territory, Jigawa, and Kano), and 3 nodes across two states (Kano and Yobe). Two additional dyadic communities were identified: one spanning Ebonyi and Bauchi, and another spanning Katsina and Kano. Across communities, Bauchi, Kano, the Federal Capital Territory (FCT), Plateau, and Yobe recur most frequently. In contrast, 181 nodes did not belong to any contagion cluster and remained isolated under this clustering criterion.

### Backbone Extraction of the Small Ruminants Mobility Network

We extracted the network backbone using a 7% threshold, which reduced the number of edges by 87% and the number of connected nodes by 84%. The resulting backbone comprised 20 nodes, the majority of which were located in Plateau State (Figure 2).

**Figure 2.**
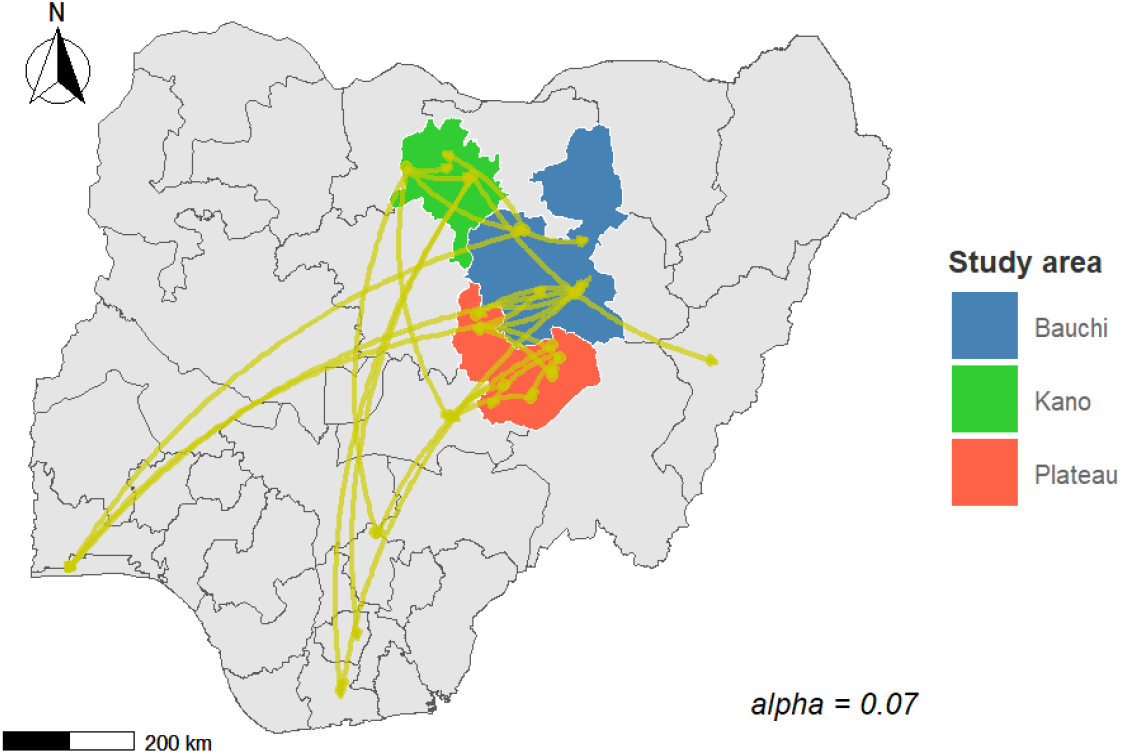
Backbone of the small ruminants mobility network. The map illustrates the backbone, extracted using a disparity filter (alpha = 0.07). 20 nodes form the network. Many of the backbone nodes are located in Plateau State (red). The nodes are dispersed from north to south. The grey areas correspond to the non-sampled States. Note that the node here represents a village.

### Impact of target Control on the Network Structure and Final Size of the Epidemic: Eliminating Backbone Nodes Drastically Reduces PPR propagation

We consider scenarios in which, upon detection of infected animals at specific nodes, movement restriction measures are implemented. These scenarios are evaluated by simulating control strategies through the following actions: (i) removing vulnerable nodes based on disease severity (Scenarios 1 to 4), (ii) eliminating the bridge links connecting previously identified contagion clusters (Scenario 5), and (iii) removing the nodes that form the network’s backbone (Scenario 6).

Overall, Figure 3 shows that interventions targeting vulnerable nodes produce only minor changes in the largest connected component (LCC), global efficiency (Eff), and total flow (TF) when compared with backbone-based intervention. Across most scenarios, the network remains highly resilient, and its overall performance is only weakly affected. The clear exception is Scenario 6, which removes backbone nodes and substantially undermines network integrity.

**Figure 3.**
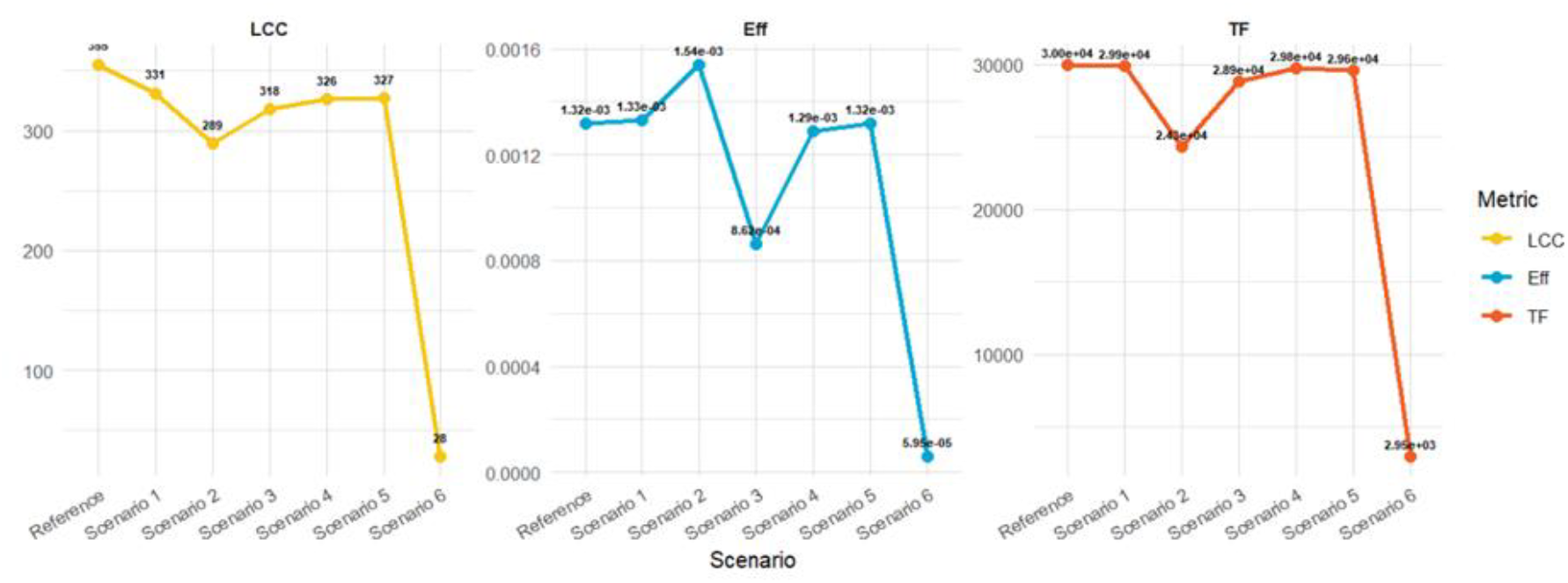
Effect of Removing Vulnerable Nodes, Bridges, and the Backbone on Network Robustness. Six network variants under the study’s scenarios: (1) Q1 vulnerable-node set removed; (2) Q2 vulnerable-node set removed; (3) Q3 vulnerable-node set removed; (4) Q4 vulnerable-node set removed; (5) inter-community bridge edges removed (“Bridges”); and (6) backbone nodes/villages removed (“Backbone”). The **Reference** is the reconstructed animal-mobility network from northeast Nigeria. Metrics shown: **LCC** = Largest Connected Component, **Eff** = global Efficiency, **TF** = Total Flow. **Q1–Q4** defined by the per-movement transmission probability (P_inf_).

Removing nodes classified at severity level 2 (Scenario 2) yields a small improvement in efficiency. In contrast, removing severity level 3 nodes (Scenario 3) provides no comparable benefit and may even slightly decrease efficiency. Likewise, although bridge links (Scenario 5) contribute to local connectivity between contagion clusters, their removal has only a marginal impact on network-wide structure and performance.

By contrast, the backbone is pivotal for maintaining both cohesion and efficiency. Removing backbone nodes collapses the LCC from 229 to 25 nodes and reduces the number of links from 355 to 28 (Figure 3), indicating severe fragmentation. Global efficiency drops from 1.32 × 10^−3^ in the reference scenario to 5.95 × 10^−5^ after backbone removal. Consistently, total movement volume through the network declines sharply, reinforcing the backbone’s central role in sustaining functional connectivity and flow

Figure 4 shows that removing vulnerable nodes in groups Q1, Q2, or Q4 does not produce a meaningful reduction in final epidemic size compared with the reference network, and eliminating bridge links yields similarly negligible effects. Targeting Q3 nodes leads to a somewhat larger reduction than these scenarios, but the impact remains modest overall. In contrast, removing backbone nodes produces by far the strongest effect, dramatically lowering the final epidemic size: under backbone control, outbreaks average approximately one infected village, regardless of the initial severity group.

**Figure 4.**
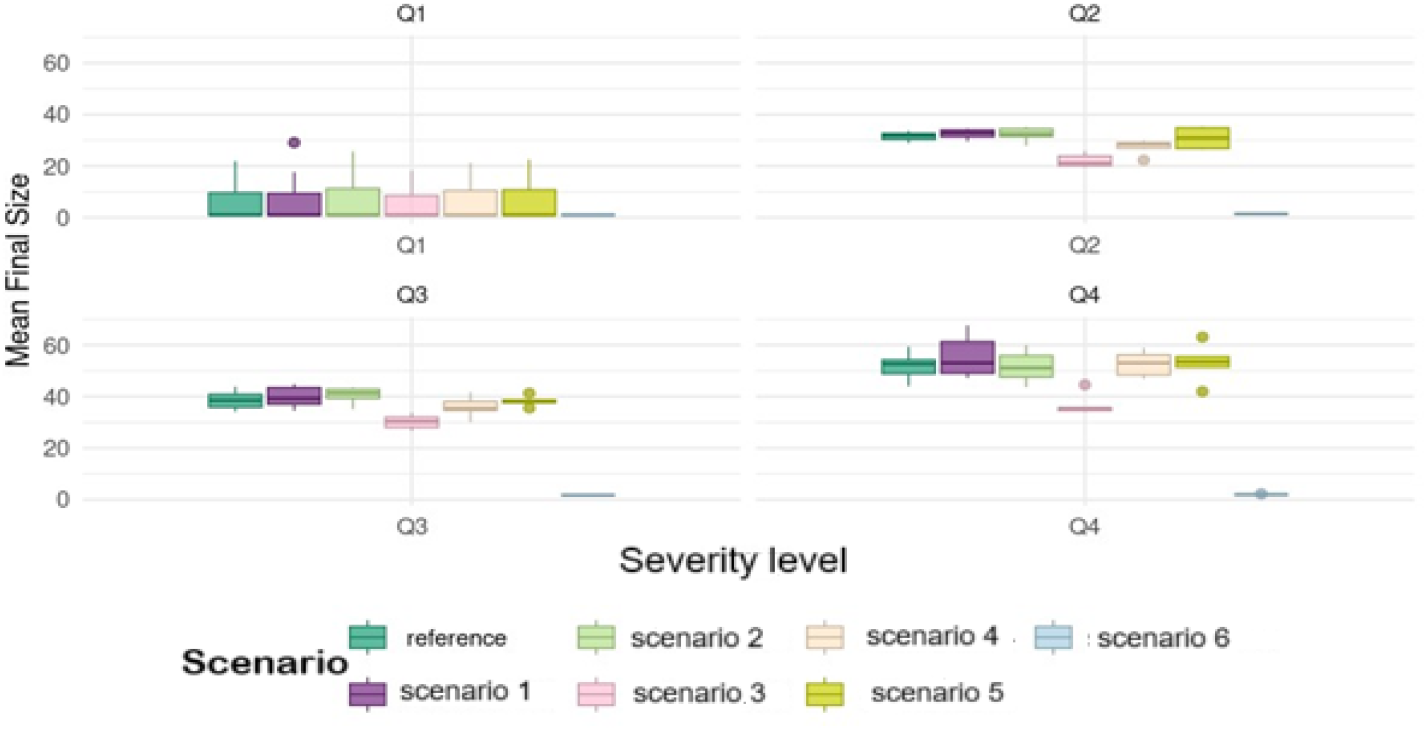
Impact of Removing Vulnerable Nodes according to the severity of the disease (from Q1 to Q2), Bridges, and Backbone on PPR Propagation: After simulating PPR using the weighted SIR model across the sixth generated network, the results were compared to the reconstructed network (reference scenario) regarding final epidemic size. Scenario 1: Network without a vulnerable node in set Q1. Scenario 2: Network without a vulnerable node in set Q2. Scenario 3: Network without a vulnerable node in set Q3. Scenario 4: Network without a vulnerable node in set Q4. Scenario 5: Network without an interlink of contagion clusters. Scenario 6: Network without a backbone.

## Discussion

In this study, we evaluated practical scenarios to strengthen PPR control, focusing on small-ruminant mobility. Given the substantial socio-economic burden of PPR in low-income settings and the limited availability of routine movement data, we prioritized strategies that are cost-effective and feasible under constrained logistical capacity.

A central result is that the network backbone represents a powerful lever for improving control of PPR-like diseases. In weighted mobility networks, backbone extraction aims to identify a parsimonious set of statistically or structurally important ties that preserves the multiscale organization of movement intensity and connectivity, while filtering redundant links (e.g., disparity-filter approaches) (Serrano et al. 2009). Our findings suggest that interventions aligned with backbone components can outperform strategies that focus on broad movement suppression, because they concentrate effort where connectivity is most epidemiologically consequential.

Beyond backbone structure, we identified contagion groups (movement communities), sets of villages with elevated mutual connectivity, whose members are likely to infect one another rapidly. Importantly, some of these groups span multiple Nigerian states, implying that once PPR is introduced into one village, spread within the group may occur quickly regardless of geographic distance. This is consistent with the general principle that community structure shapes both epidemic dynamics and the comparative efficiency of targeting strategies. As shown by Salathé et al. (2010) (Salathé and Jones 2010), in networks with strong community structure, immunization focused on bridging individuals that connect communities can be more effective than targeting only the most highly connected individuals. Related work similarly emphasizes “bridge hubs” and community-aware targeting as efficient control heuristics when resources are limited (Gong et al. 2013). Operationally, this supports a vaccination and movement-management logic that prioritizes backbone-informed nodes/edges and community “gateways” that facilitate between-group spread. Importantly, this approach does not imply blanket movement restrictions across the livestock mobility network. Rather, it focuses surveillance, vaccination, and movement-control efforts on a limited number of strategically positioned villages and connections that disproportionately contribute to disease transmission.

In contrast, our results indicate that targeting inter-community links alone is less relevant or insufficient to prevent spread across the full mobility network. Large-scale prohibitions on movement along major trade routes may be inefficient and can trigger rerouting or non-compliance, potentially shifting rather than eliminating risk. This aligns with broader PPR control discussions highlighting why movement restrictions are difficult to sustain in many production systems where markets, feed access, and seasonal mobility are fundamental (Cameron 2019). Targeted interventions, rather than blanket bans, have also been shown in other livestock contexts to outperform random strategies and to be more robust under operational constraints (Gates and Woolhouse 2015; Chaters et al. 2019). Where enforcement is considered, random checkpoints along major routes may be conceptually appealing but often face feasibility and safety limitations; a more practical alternative is to monitor gateway locations that are locally central yet strategically positioned to mediate movements between communities. Such “sentinel” or “chokepoint” logic is also coherent with network-informed surveillance approaches in livestock systems (Chaters et al. 2019).

Several limitations should be acknowledged. First, we modeled a static network, whereas real livestock mobility is time-varying, shaped by seasonality, markets, insecurity, and shocks (Holme and Saramäki 2012). Temporal fluctuations can substantially affect which nodes and links appear most influential at a given time, and outbreak outcomes may be sensitive to initial conditions (Bajardi et al. 2012). Second, backbone membership itself can evolve as mobility patterns shift. Temporal and inference-based extensions for identifying “irreducible” or time-robust backbone structure have been proposed (Nadini et al. 2020) and could improve targeting when longitudinal movement data become available. Third, movement networks in endemic settings are frequently reconstructed from incomplete or biased samples rather than continuous traceability systems, which can affect estimated centralities and community structure; methods to address fragmentation and inference uncertainty remain an active area (Leung et al. 2023).

Despite these constraints, our results provide a concrete operational implication: backbone-informed targeting offers a plausible path to increase the efficiency of control and vaccination under limited resources. This complements the FAO/WOAH emphasis on risk-based, progressive strengthening of surveillance and vaccination systems, including the role of animal identification and movement management where feasible (FAO/WOAH PPR GEP; PPR Global Strategy). Field evaluation, assessing feasibility, acceptability, and measured epidemiological impact, will be essential to translate these network insights into demonstrable control gains.

## Conclusion

In conclusion, our results show that integrating animal-movement information into network-based targeting can improve the efficiency of PPR control. In particular, prioritizing backbone villages, and the community “gateways” they reveal, offers a pragmatic way to concentrate limited resources where they are most likely to disrupt transmission, which is especially relevant in low-income settings with constrained vaccination capacity and incomplete movement tracking. More broadly, the study supports the value of adaptive, risk-based control strategies that exploits network structure rather than relying on uniform coverage. Future work should extend these findings using time-varying movement data, broader geographic coverage, and field validation to assess feasibility, compliance, and measurable epidemiological impact across diverse production and mobility contexts.

## Notation

Conceived and designed the analysis: AA, MA, AM, SE; Collected and curated the data: SI, MB; Performed the analysis: AM; Wrote the draft paper: AM; Critical review: AA, MA, MC, EC.

## Data, Scripts, and Code Availability

A total of 1,065 stakeholders (traders, transporters, and farmers) were interviewed in 10 Local Government Areas (LGAs) between April 1, 2022, and September 30, 2022. A convenience sampling approach was used, with participants recruited voluntarily while attending livestock markets. Respondents provided information on the origin and destination of animal movements (country, state, LGA, and village), the species transported (goats, sheep, or both), the number of animals moved, the frequency of market visits, and the purpose of the movement. These data were used to construct livestock mobility networks at several administrative levels (village, district, and LGA), with edge weights corresponding to the number of animals moved.

The movement dataset used in this study is publicly available on Zenodo: https://zenodo.org/records/13121488 and includes origin–destination movement records aggregated at the administrative district level.

The scripts used for data processing, network analysis, epidemic simulations, and identification of vulnerable (sentinel) nodes are available from the Zenodo: https://doi.org/10.5281/zenodo.20802295

The repository includes scripts for:

- descriptive analysis of livestock movement data;
- construction and analysis of livestock mobility networks;
- simulation of PPR-like disease diffusion using stochastic SI and SIR network models;
- identification of vulnerable (sentinel) nodes;

Contagion clusters were identified using the COCLEA algorithm (Nath et al., 2019). The implementation used in this study was provided by the original authors and was not modified.

Network backbone extraction was performed using the disparity-filter method (Serrano et al., 2009) implemented in publicly available software packages. No modification of the original algorithm was introduced.

## Funding

The research was funded by a grant from European Commission (Development Cooperation Instruments) awarded to the project ‘EU

Support to Livestock Disease Surveillance Knowledge Integration -LIDISKI’ (FOOD/2019/410-957).

## Footnotes

1 Lidiski | Livestock Disease Surveillance Knowledge Integration

